# Embryonic geometry underlies phenotypic variation in decanalized conditions

**DOI:** 10.1101/579623

**Authors:** A. Huang, T. E. Saunders

## Abstract

During development, many mutations cause increased variation in phenotypic outcomes, a phenomenon termed decanalization. Such variations can often be attributed to genetic and environmental perturbations. However, phenotypic discordance remains even in isogenic model organisms raised in homogeneous environments. To understand the mechanisms underlying phenotypic variation, we used as a model the highly precise anterior-posterior (AP) patterning of the early *Drosophila* embryo. We decanalized the system by depleting the maternal *bcd* product and found that in contrast to the highly scaled patterning in the wild-type, the segmentation gene boundaries shift away from the scaled positions according to the total embryonic length. Embryonic geometry is hence a key factor predetermining patterning outcomes in such decanalized conditions. Embryonic geometry was also found to predict individual patterning outcomes under *bcd* overexpression, another decanalizing condition. Further analysis of the gene regulatory network acting downstream of the morphogen identified vulnerable points in the networks due to limitations in the available physical space.

## Introduction

The phenomenon of canalization describes the constancy in developmental outcomes between different individuals within a wild-type species growing in their native environments [1–3]. To better understand how canalized a developmental process is, we need to quantitatively measure the molecular profiles of the developmental regulatory genes in multi-cellular organisms. In *Drosophila,* the highly reproducible body patterning in adult flies originates from both the reproducible setup of the instructive morphogen gradients and the precise downstream transcriptional readouts early in embryogenesis [4–6]. In particular, the inter-individual variation of the positional information conferred by gene expression is below the width of a single cell [4,7]. This means that these developmental processes are highly canalized. Given the ubiquity of canalization in nature, such highly reproducible developmental processes are likely not exclusive to insect development.

The developmental canalization that we see in contemporary species is the product of evolution, either as the consequence of stabilizing selection [8] or the manifestation of the intrinsic properties of the underlying complex gene regulatory networks [9]. Canalization can break down in individuals subjected to aberrant genetic mutations or extreme environmental conditions [10,11]. Such individuals in decanalized conditions become sensitive to variations in their genetic background and external environments, which are otherwise neutral to developmental outcomes. This leads to significantly increased inter-individual variation in phenotypic outcomes.

It is important to characterize the sources of variation in order to understand what canalization is actually buffering against. Interestingly, significant phenotypic variation remains in laboratory animals with isogenic genomes, exposed to homogeneous environments [12]. This indicates that other components besides genetic and environmental variation cause phenotypic discordance under decanalized conditions. Previous work has proposed that stochastic expression of redundant genes predicts the developmental outcome of the mutant individuals [13]. However, in many other cases, it remains elusive as to why mutation increases inter-individual phenotypic variation and what alternative components underlie such variation [14,15].

Therefore, to identify the potential sources of variation that govern phenotypic variation, we utilized early *Drosophila* embryonic patterning. The developmental process was decanalized by either knocking out the maternal *bicoid (bcd)* gene or introducing aberrantly high amplitude of the Bcd morphogen gradient, both resulting in increased inter-individual variation of patterning outcomes [16,17]. We found that the naturally variable embryonic geometry acts as a previously unidentified source of variation that predetermines the individual phenotype. Further, we found that specific segmentation identities are preferentially affected in decanalized conditions, which is informative of the epigenetic interactions of the underlying developmental genes.

## Results

### Lack of maternal bcd results in increased variation in patterning outcomes

The cuticle phenotype of maternal *bcd* mutants was previously characterized using a series of different *bcd* alleles, with increasing allelic strength showing defects reaching further into posterior regions of the embryonic patterning [16,18]. Phenotypic variation was also observed among embryos derived from females carrying the same mutant allele [16,18]. For example, the amorphic *bcd^E1^* allele gives rise to embryonic patterning lacking the head and thoracic structures with 100% penetrance and replaced by duplicated posterior spiracles. Comparatively, the patterning outcomes ranging from abdominal segments 1 (A1) to A5 are highly variable among different individuals, manifesting in either fusion or depletion of various number of denticle belts. Meanwhile, segments posterior to A6 (inclusive) always remain intact.

To systematically understand the inter-individual phenotypic variation among *bcd* mutants, we utilized an allele generated by the CRISPR-MiMIC method [19,20]. The MiMIC transposon carrying stop codons in all three reading frames is targeted by CRISPR to insert into the first intron of the endogenous *bcd* gene. Therefore, no functional Bcd protein is produced by this knockout allele (annotated *bcd^KO^*). Moreover, the MiMIC construct contains an eGFP reporter, which facilitates further genetic manipulation, such as recombination, carried out in this study.

The cuticular pattern of the *bcd^KO^* allele qualitatively recapitulates that of *bcd^E1^* (Fig 1A-F, Fig S1A). While the anterior embryonic patterning is entirely defective, the patterning defects in posterior regions are more variable. The number of normal abdominal denticle belts in each embryo ranges from three (A6 – A8) to seven (A2 – A8), with four intact abdominal segments (A5 –A8) being the most frequently observed phenotype (Fig 1G). Structures indicating partially differentiated abdominal segments can be observed in the anterior regions of the *bcd^KO^* embryos. However, these structures do not recapitulate any of the wild-type denticle belts (Fig 1A-F). Further, we observed and classified the variable phenotypes of the duplicated posterior spiracles according to the completeness of the organ morphogenesis (Fig 1H-I). Interestingly, the fully developed ectopic spiracle organ (Fig 1H(i)) is only observed in embryos showing less than six intact abdominal segments, while most frequently observed in individuals with three intact abdominal segments (Fig 1G, dashed bar area). This indicates correlation between the patterning outcomes of the abdominal regions and the ectopic spiracles.

**Fig 1.**
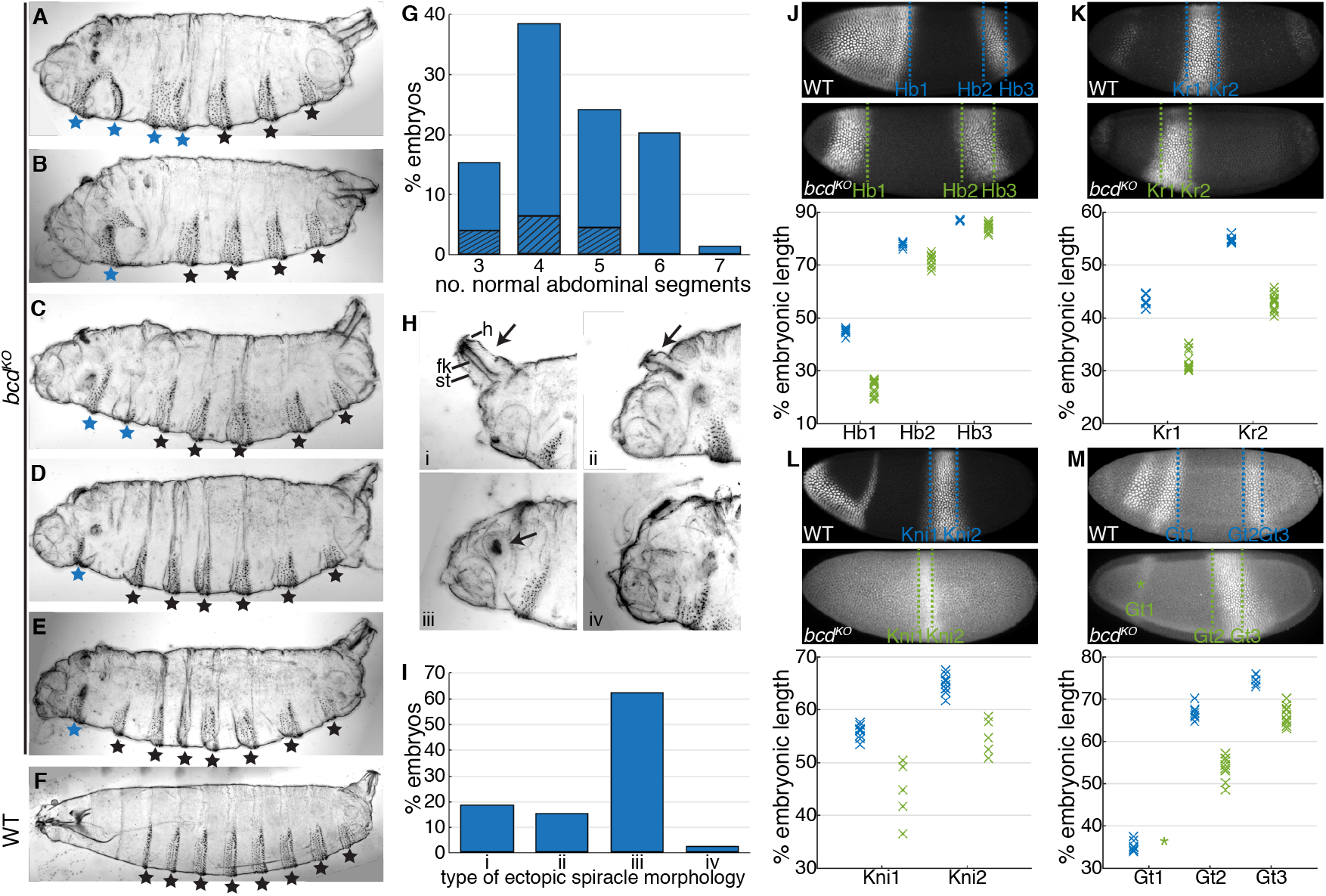
*bcd* mutants show increased variation in cuticles and gap gene expression patterns. (A-F) Representative cuticle patterns of maternal *bcd^KO^* mutant individuals (A-E) and wild type (F). Black stars indicate normal abdominal segments; blue stars indicate aberrant denticle belts. (G) Phenotypic frequency showing different number of normal abdominal segments in *bcd^KO^* mutant individuals (n = 202). Dashed area indicates proportion of embryos showing fully developed ectopic spiracles as shown in H(i). (H) Various phenotypes of ectopic posterior spiracle observed in the anterior region of the *bcd^KO^* embryos. Arrows point to ectopic spiracles showing complete morphogenesis (i), partial morphogenesis (ii), only primordial structures (iii), and no ectopic spiracle differentiation (iv). Letter h indicates spiracular hair; fk, filzkörper and st, stigmatophore. (I) Phenotypic frequency of different ectopic spiracle morphology as shown in H (n = 168). (J-M) Gap gene expression by the end of blastoderm stage in wild type (upper panel) and *bcd^KO^* (lower panel). Dashed lines mark the boundary of expression domains, and the scaled boundary positions of individuals are plotted in the graphs below (wild type in blue and *bcd^KO^* in green). Asterisk indicates weak transcriptional activation of anterior Gt domain.

The wide spectrum of *bcd^KO^* cuticle phenotypes can be attributed to the variation of patterning gene expression during the blastoderm stage. In the absence of a maternal Bcd gradient, the antero-posterior (AP) axis of the embryos is patterned by the remnant maternal positional cues including: the anterior Hunchback gradient; the uniformly distributed Caudal; the posteriorly localized Nanos; and the Torso pathway activated at both terminal regions [21,22]. By the end of the blastoderm stage, the relative boundary positions of all gap gene expression domains show significant anterior shift compared to those in the wild-type embryos (Fig 1J-M). Importantly, in the wild-type condition, the inter-individual variation of most boundary positions is at the range of 1% embryonic length (EL), with an exception being the posterior boundaries of Giant (Gt) and Knirps (Kni), which have positional variation slightly over 1.5 % EL [7]. In comparison, the absence of Bcd activity results in significantly increased variation in gap gene boundary positions (Fig S1B, p<0.01 for all measured boundaries to have increased error randomly), with the anterior Kni boundary being the most variable, showing variation above 5% EL (Fig 1J-M). Occasionally we detected no Krüppel (Kr) nuclear intensity in the presumed expression region in *bcd^KO^* individuals (2 out of 15 individuals), indicating the failure of Kr gene activation in these embryos (Fig S1C). The anterior Gt domain shows similar inter-individual variation, with the majority of the embryos failing to properly activate anterior Gt expression (8 out of 10 individuals). Instead only a thin stripe of diminished cytoplasmic signal can be detected (Fig 1M, asterisk; Fig S1D). It is noteworthy that embryos derived from a single pair of *bcd^KO^* parents raised in constant environmental conditions show equivalent phenotypic variation (Fig S1E and F), suggesting that the inter-individual variation observed cannot be attributed to differences in either environment or genetic background.

### AP Patterning of bcd mutants correlates with embryonic length

We next wanted to understand the causes of reduced developmental reproducibility in the absence of Bcd-instructed positional information and identify potential sources of variation that predetermine patterning outcomes of each mutant individual. We monitored the dynamic expression of the segment-polarity gene *engrailed* (*en*) in different *bcd^KO^* individuals. All the control embryos show invariably eleven En stripes, demarcating the posterior boundary of each body segment including three thoracic (T1~T3) and eight abdominal segments (A1~A8; Fig 2B). In contrast, different numbers of En stripes are activated in *bcd^KO^* embryos, consistent with the variable number of intact denticle belts that we observed previously. This number remains unchanged throughout embryogenesis. We also measured the geometrical parameters of each embryo (Fig 2A). Interestingly, we found that the number of En stripes positively correlates with the embryonic length along the AP axis (Fig 2C-G, Video 1). These results led us to hypothesize that, in the absence of maternal *bcd* activity, the number of body segments generated during pattern formation is dependent on the embryo absolute length. Therefore, embryonic geometry could be a previously unrecognized source of variation that underlies phenotypic variation in patterning outcomes of mutant individuals.

**Fig 2.**
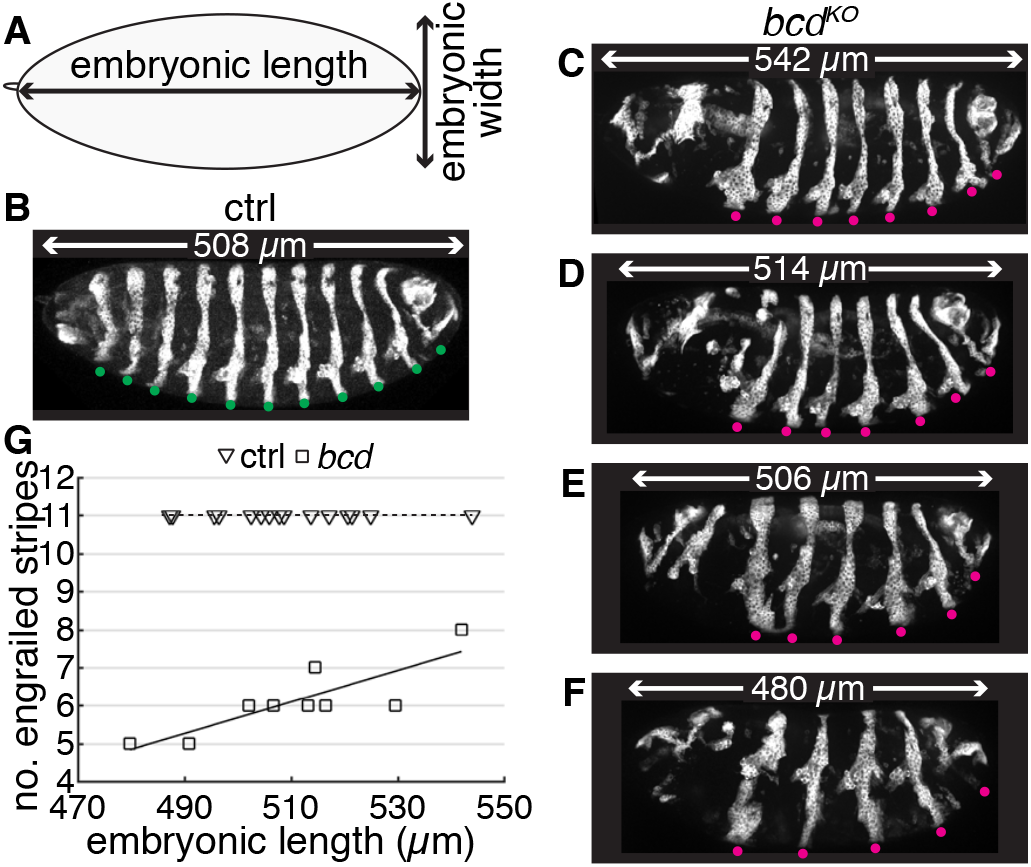
*bcd* mutant phenotypic outcomes correlate with embryonic length. (A) Graphic demonstration of parameters of embryonic geometry. Embryonic length refers to the distance between anterior and posterior poles, and embryonic width refers to the widest dorso-ventral distance along AP axis. (B-F) En expression at the end of head involution stage in ctrl (B) and *bcd^KO^* (C-F) embryos. Dots indicate En stripes (ctrl, green; *bcd^KO^*, magenta)and the numbers indicate embryonic length.

### Developmental reproducibility is preserved in a wide range of embryonic geometry

To test our hypothesis, we first asked if altering embryonic geometry alone can alter embryonic pattern formation. The geometry of each embryo is predetermined during oogenesis when the follicle cells surrounding the egg chamber transform the developing egg from a sphere to an ellipse [23,24]. This process is mediated by the planar cell polarity (PCP) of the follicle cells and the elliptical shape of the embryos remains unchanged throughout embryogenesis. Here, we used maternal ShRNA to knockdown one of the PCP core components, atypical cadherin Fat2 in the follicle cells [25], and we henceforth refer to these as *fat2RNAi* embryos. This reduces the embryonic length from 510 ± 17 (s.d.) μm in wild type to 432 ± 40 μm in *fat2RNAi* embryos (Fig 3A). Meanwhile, the perturbed embryos show an increased embryonic width (EW) along the dorso-ventral axis (Fig S2A and B, 196 ± 5 μm in wild type and 202 ± 11 μm in *fat2RNAi*). Together, these geometrical variations lead to only a slight reduction ~8 % in the embryonic size (assuming an ellipsoidal geometry) compared to wild type embryos (Fig 3B). Depletion of the *bcd* gene product does not affect embryonic geometry (Fig 3A and B). The round eggs of *fat2RNAi* embryos are fertilizable and continue with proper embryogenesis (Video 2).

**Fig 3.**
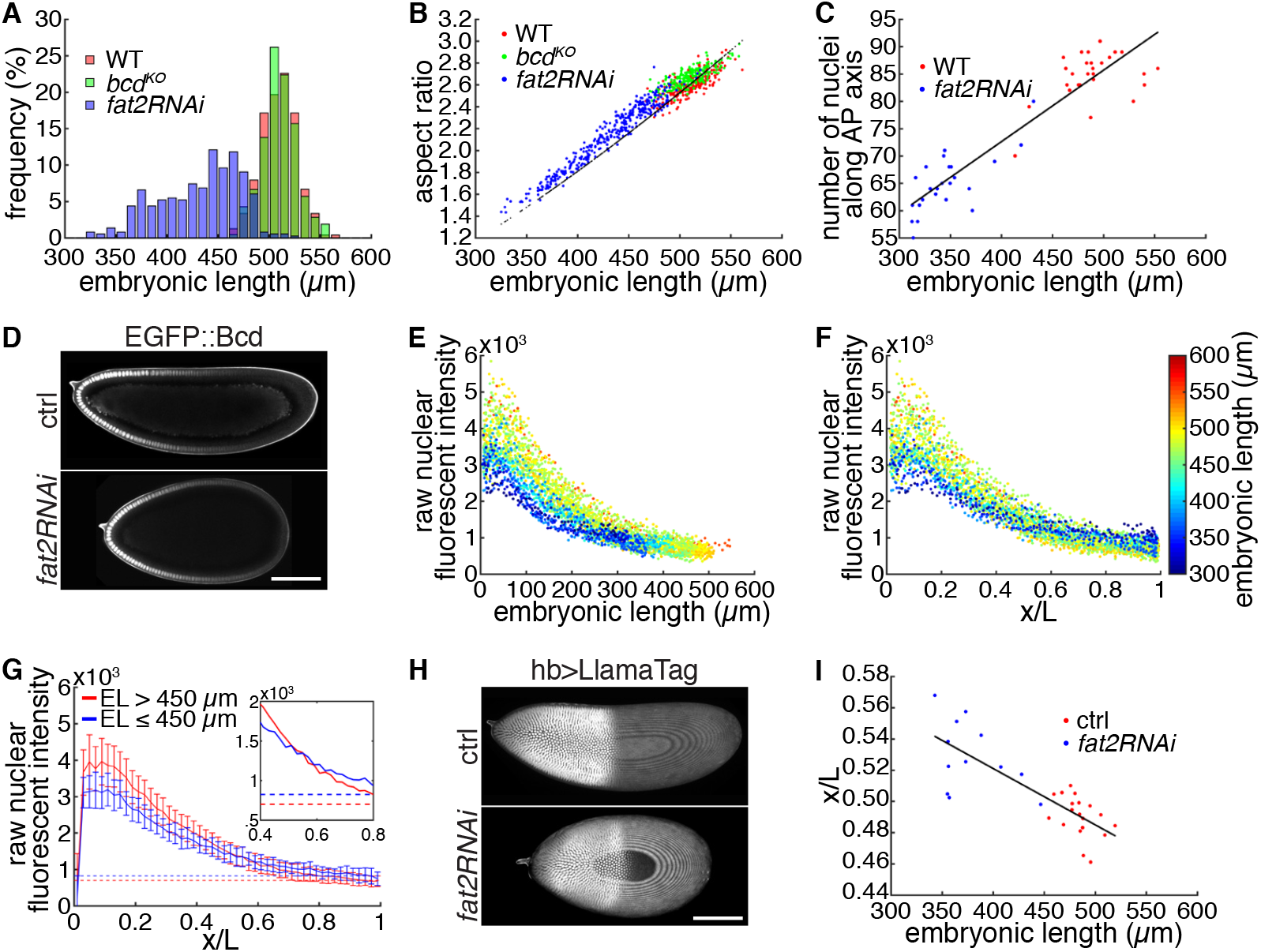
Embryonic patterning in response to perturbed embryonic geometry. (A) Distribution of embryonic length in wild type (n = 239), maternal *bcd^KO^* (n = 210) and *fat2RNAi* (n = 364) embryos. (B) Aspect ratio (EL/EW) is plotted against embryonic length for each individual embryo in three genotypes. The black dots indicate the expected aspect ratio given the measured embryonic length in each individual if embryonic size is conserved as the mean of the wild-type. (C) Number of nuclei along the AP axis is plotted against embryonic length in wild type and *fat2RNAi* embryos. Line indicates linear regression of all data. (D) Midsagittal plane of ctrl (top) and *fat2RNAi* (bottom) embryos expressing eGFP::Bcd in mid n.c. 14. (E-F) EGFP::Bcd profiles of both ctrl and *fat2RNAi* embryos plotted in a single graph as a function of absolute distance from the anterior pole (E) or scaled AP position (F). Each dot represents the average concentration in a single nucleus. Colormap indicates the absolute AP length of each individual. (G) Mean and standard deviation of nuclear intensity within each 2% EL were computed for group of embryos longer (red, n = 27) and shorter (blue, n = 17) than 450 μm. Dashed lines indicates the mean intensity in the most posterior 10% of the EL. Inset is the zoomed plot without errorbars to show the intersect of two curves. (H) Max projection of ctrl (top) and *fat2RNAi* (bottom) embryos expressing maternally loaded eGFP and hb>LlamaTag. (I) Scaled Hb boundary positions plotted against EL in two genotypes. Scale bar, 100μm.

We examined the nuclear distribution in the blastoderm embryos, as nuclei are the basic units interpreting positional information, and an altered nuclear distribution may affect patterning outcomes. We found that nuclear number along the AP axis decreases proportionally to embryonic length, leaving the inter-nuclear distance unchanged (Fig 3C; Fig S2C). In other words, the number of nuclei to interpret AP positional information reduces from 85 ± 4 in wild type to 65 ± 5 in *fat2RNAi embryos*.

Next, we investigated how establishment of the Bcd morphogen gradient is influenced by the altered embryonic geometry in *fat2RNAi* embryos. We live imaged eGFP-Bcd fusion protein in both control and *fat2RNAi* backgrounds, and measured the nuclear Bcd intensity along the AP axis at mid nuclear cycle (n.c.) 14 (Fig 3D). We found that the absolute Bcd concentration is lower in the anterior half of the embryo in *fat2RNAi* individuals compared to that of control (Fig 3E-F). Further, Bcd profiles from different individuals intersect near the mid region of the embryo, with shorter embryos showing higher Bcd concentration in the posterior region (Fig 3G). These observed Bcd gradient profiles in different embryonic geometry are consistent with the SDD model [26] of Bcd gradient formation in different geometries (Methods and Fig S2D).

We predicted that the ‘flattening’ of the Bcd profile in response to decreasing aspect ratio of the embryo should result in a slight but measurable shift in the downstream gap gene expression domains. We first measured Hunchback (Hb) expression boundary at mid n.c. 14 using live imaging of *hb>LlamaTag* [27]. In control embryos, the Hb expression boundary locates at 49.0% EL with variation of 1.3% EL, consistent with previous reports [28]. Comparatively, the Hb boundary shows a posterization in *fat2RNAi* embryos (52.9% EL) with an increased variation of 2.3% EL (Fig 3H and I). However, considering the absolute length of *fat2RNAi* embryos, such variation indicates that the Hb boundary is still defined at the precision of a single nucleus domain. Other gap genes show similar posterization and variation in their boundary positions (Fig S2E-H).

These results suggest that, when we manipulate the embryonic geometry to an extent beyond that naturally observed, the reproducibility of the patterning outcomes is preserved as compared to that of the wild-type scenario. Therefore, the intact early embryonic patterning network is highly robust to variations in embryonic geometry.

### Embryonic length dictates segmentation gene pattern in the absence of bcd

We now return to our hypothesis that embryonic geometry is a potential source of variation that predetermines phenotypic outcomes of mutant individuals. We introduced the *fat2RNAi* knockdown into a *bcd^KO^* background to see how embryonic patterning is affected accordingly. The cuticle pattern of embryos derived from *fat2RNAi, bcd^KO^* females resembles that of *bcd^KO^* alone, but the number of properly patterned abdominal denticle belts reduces with decreasing embryonic length. Moreover, novel phenotypes showing only one or two abdominal segments were observed when the embryonic length drops beyond the natural range (Fig 4A and B). All of the *fat2RNAi, bcd^KO^* embryos showed duplicated spiracles with fully developed morphology (see Fig 1H(i)), consistent with our previous result that such spiracles prevail in embryos with shorter AP length. Similar observation was made with En expression. The number of En stripes shows positive linear correlation with embryonic length in *bcd^KO^* embryos (R^2^ = 0.925), while individuals with only *fat2RNAi* knockdown express exclusively eleven En stripes regardless of their embryonic length (Fig 4C-E). We noticed that *fat2RNAi* embryos shorter than 400 μm developed morphological defects in late embryogenesis, where abnormal dorsal closure leads to mismatch between the two lateral sides of the ectoderm (Fig 4D, arrows). However, such local morphological abnormality is likely due to defective tissue morphogenesis as a consequence of limited physical space, rather than patterning errors (Video 3).

**Fig 4.**
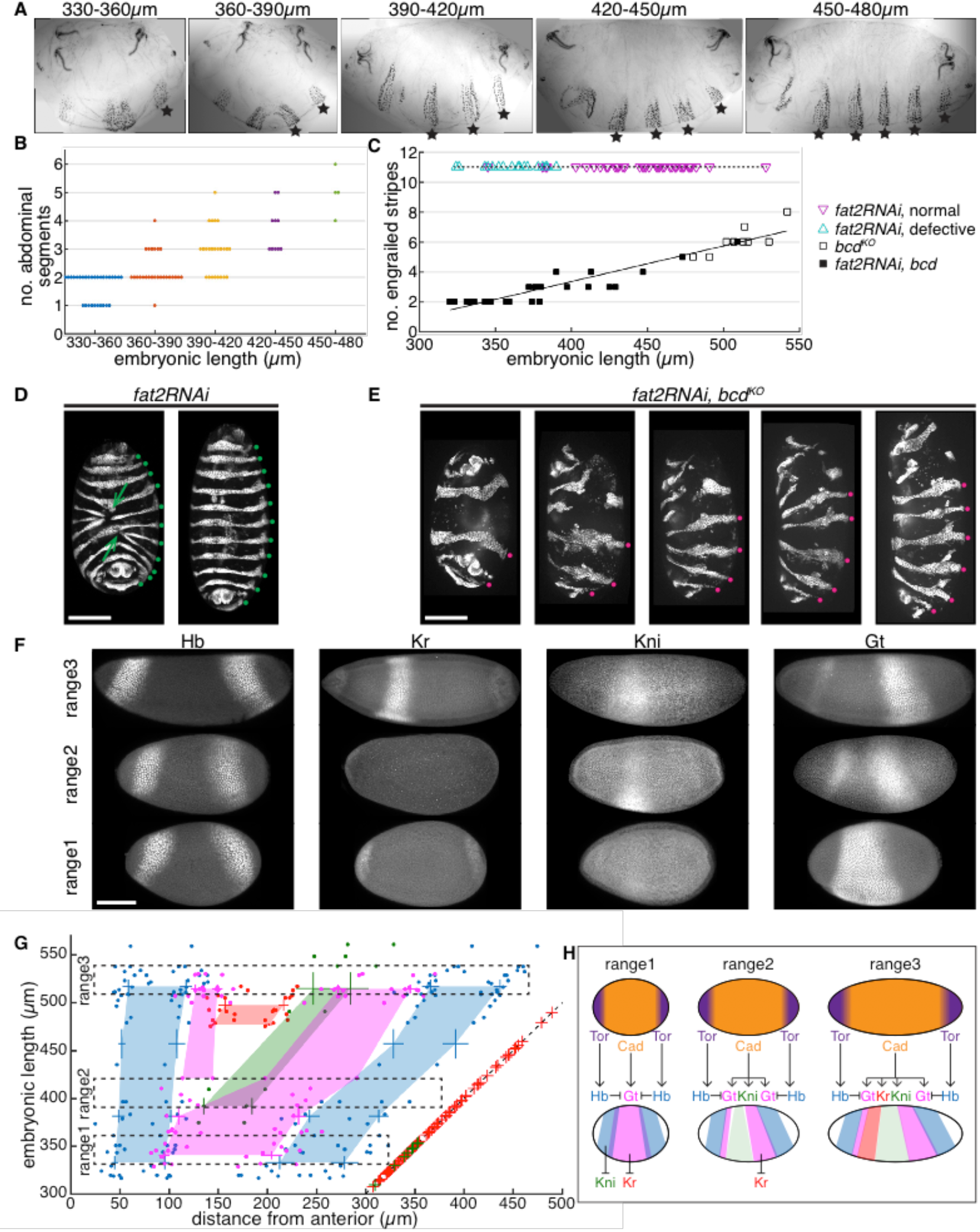
Embryonic geometry underlies patterning outcomes of *bed^KO^* individuals. (A) Representative cuticle patterns of *fat2RNAi, bcd^KO^* embryos within different ranges of embryonic length, from left to right, 330-360, 360-390, 390-420,420-450, and 450-480μm. Stars indicate normal abdominal segments. (B) Number of normal abdominal segments plotted against EL range of each individual. (C) Number of En stripes plotted against EL in individuals from three genotypes. Magenta triangles indicate individuals showing defective morphogenesis at the end of dorsal closure; cyan triangles indicate normal morphogenesis. (D) En expression in *fat2RNAi* embryos showing defective dorsal closure (left) or normal morphogenesis (right). Arrows indicate locations of defects. Green dots indicate En stripes. (E) Representative images of *fat2RNAi, bcd^KO^* embryos showing different number of En stripes. Magenta dots indicate En stripes. (F) Representative gap gene expression patterns within different range of EL. Range1, 330-360μm; range2, 390-420μm; and range3, 510-540μm. (G) Embryonic length of *bcd^KO^* individual is plotted vs. the boundary position of four gap genes shown as absolute distance from the anterior pole (Hb, blue; Gt, magenta; Kr, red and Kni, green). Data from every 30μm EL interval were binned to compute mean and s.d. and the colored areas are generated by connecting mean values of different EL ranges. Dashed boxes indicate ranges of EL corresponding to (F). Dashed line indicates posterior boundary of individual embryos; red and green crosses overlapping the dashed line indicate individuals with corresponding EL not expressing Kr and Kni, respectively. (H) Schematic illustration of positional information transfer from maternal systems to gap gene expression in *bcd^KO^* embryos within different range of embryonic length. Scale bar, 100μm.

A longstanding question in patterning is how do gene regulatory networks downstream of morphogens incorporate information about the macroscopic geometrical parameters of each individual to give rise to scaled patterning outputs? We can begin to tackle this question by asking how the gene expression boundaries vary with embryonic geometry in *bcd^KO^* mutants. Fig 4F shows representative expression patterns of four gap genes in *bcd^KO^* mutants. The gene network shows qualitative differences in behavior within different ranges of embryonic length. Without the long-range gradient of Bcd, zygotic *hb* transcription is activated by the termini system mediated by the terminal gap gene, *tailless* [29](Fig S3A). As a result, two Hb stripes form near the anterior and posterior poles of the embryo, spanning a width of ~10 and 15 nuclei, respectively. In embryos with extremely large aspect ratio (range 1, EL within 330 – 360 μm), the two Hb expression domains are in close proximity. This inhibits the expression of Kni, which is strongly repressed by Hb, in the central region of the embryo [30]. Meanwhile, Gt is activated by uniformly distributed maternal Cad protein, and in turn inhibits Kr expression [31,32](Fig 4F-H, range 1).

In individuals with increased embryonic length (range 2, EL within 390 – 420 μm), Hb stripes in the terminal regions separate further apart, permitting Kni expression in the middle region (Fig 4F-H, range 2). This Kni stripe is sandwiched by two Gt expression domains, a thin anterior stripe and a wider posterior one. The anterior Gt stripe in *bcd* mutants has been observed before [33] but its regulatory interactions remain elusive. Potentially it is activated by the remnant anterior determinants such as the maternal Hb, distributed in the anterior half of the embryo [22]. Comparatively shorter embryos show phenotypically higher degree of symmetry in both cuticle and gene expression patterns, conceivably due to stronger repression of maternal Hb in shorter individuals (Fig S3B-D).

Looking more closely at individuals within the natural range of embryonic geometry (range 3, EL within 510 – 540 μm), the sufficient physical space between two Hb stripes permits the expression of Kr, Kni and Gt, arranged in spatial order that is conserved as in wild-type embryos (Fig 4 F-H, range 3). In summary, as a consequence of gradually increasing embryonic length, a continuously increasing variety of gap gene expression domains are activated along the AP axis, which is in turn, translated into increased number of body segments, as manifested by the pair-rule gene expression pattern (Fig S3E).

### The gene network breaks at susceptible point in decanalized conditions

Phenotypic discordance has been previously observed as a consequence of artificially altered maternal *bcd* dosage, with an increasing number of *bcd* gene insertions leading to a larger proportion of individuals showing defective patterning [17]. Hence, we wanted to know if embryonic geometry also underlies inter-individual variation under decanalized conditions of *bcd* overexpression. To effectively increase the Bcd gradient amplitude, we generated a tandem *bcd* construct, where two copies of the *bcd* gene are linked by the P2A self-cleaving peptide (Fig 5A). Two transgenic fly lines with two and four genomic insertions of this construct deposit *bcd* mRNA into embryos at ~3 (6x *bcd*) and ~5 (10x *bcd*) fold that of the wild-type amount, respectively (Fig 5B; Fig S4A). As Bcd protein counts scale linearly with that of its mRNA, the corresponding amplitude of the Bcd gradient are expected to show the same fold changes [6], as manifested by the posterior displacement of cephalic furrow position (Fig 5B).

**Fig 5.**
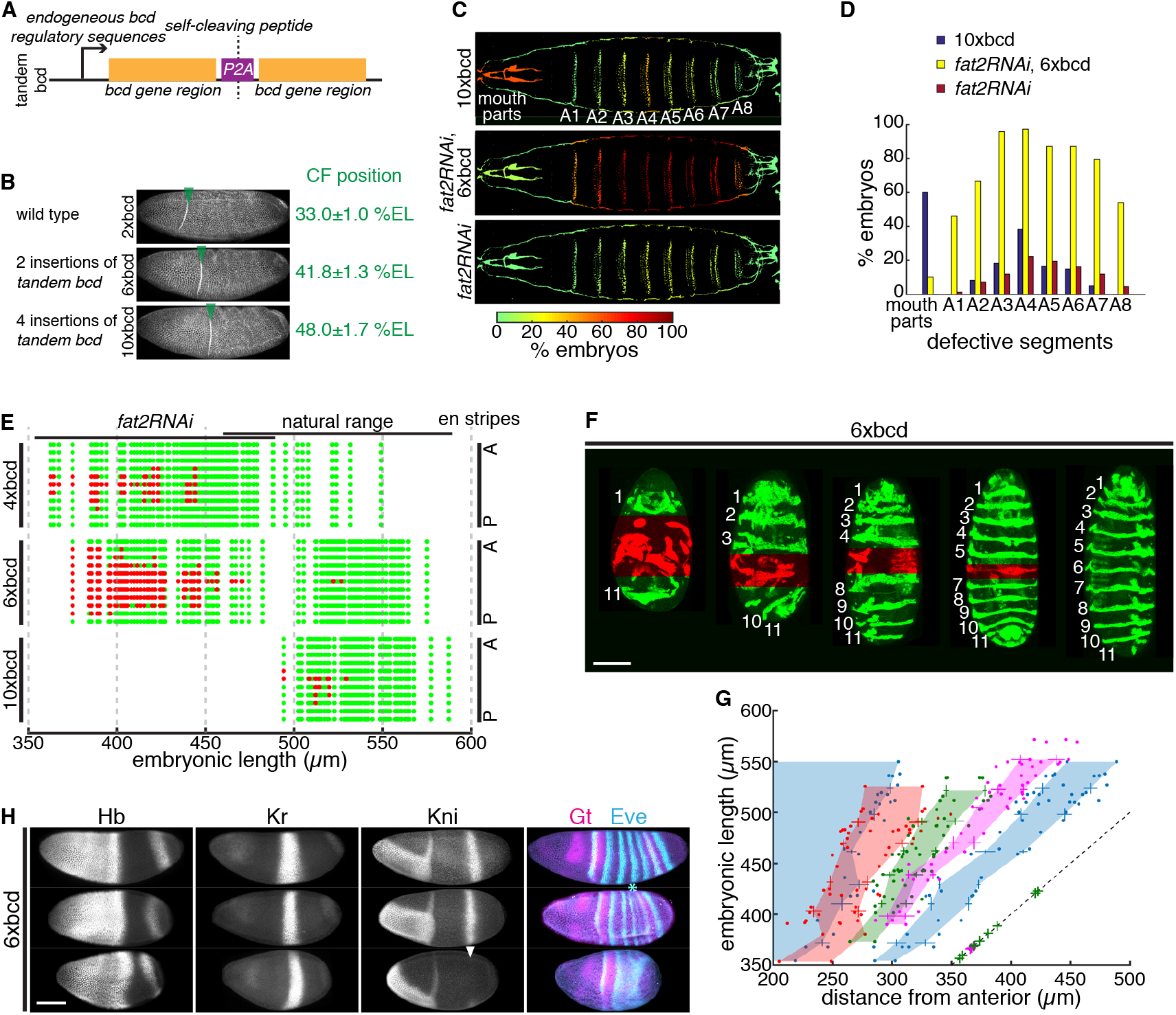
Embryonic patterning breaks down at A4 segment with *bcd* overexpression. (A) Schematic illustration of tandem *bcd* construct. (B) Embryos expressing 2x (wild type), 6x and 10x of maternal *bcd* fixed at the onset of gastrulation and stained with Phalloidin. Green arrowheads indicate the positions of cephalic furrow (CF) formation. (C) Distribution of defective cuticular segments in non-hatched 10x*bcd* (top), *fat2RNAi,* 6x*bcd* (mid) and *fat2RNAi* (bottom) embryos. Colormap indicates the frequency of defects in each segmental region. (D) Bar plots showing the distribution of defective cuticular segments in three genotypes. (E) En expression patterns in 4x (top), 6x (mid) and 10x (bottom) *bcd* embryos with different EL. Green dots, normal En stripes; red dots, defective En stripes; A, anterior; P, posterior. (F) Representative 6x*bcd* embryos with different EL expressing *en>mCD8::GFP.* Numbers indicate En stripe identities and red mask indicate defective segmental regions. (G) Embryonic length of 6x*bcd* individual is plotted vs. the boundary position of four gap genes shown as absolute distance from the anterior pole (Hb, blue; Gt, magenta; Kr, red and Kni, green). Data from every 30μm EL interval were binned to compute mean and s.d. and the colored areas are generated by connecting mean values of different EL ranges. Dashed line indicates posterior boundary of individuals; green and magenta crosses overlapping the dashed line indicate individuals with corresponding EL not expressing Kni and Gt, respectively. (H) Representative segmentation gene expression in 6x*bcd* embryos with different EL. Asterisk indicates repressed Eve stripe 5 and arrowhead indicates failed activation of Kni. Scale bar, 100μm.

The ~5-fold *bcd* overexpression compromises viability to adulthood and the non-hatched embryos displayed a plethora of defective patterning phenotypes (Fig 5C; Fig S4B). Individuals with mild defects frequently displayed missing or fused denticle belts in A4 segment (Fig 5C-D), a positional bias that has been reported previously [17]. More severe phenotypes showed defects in a spreading region centered about the A4 segment. Meanwhile, embryos show high rate of mouth defects as a consequence of significantly increased local Bcd concentration in the most anterior region (Fig 5CD; Fig S4C). Patterning defects were rarely seen in 6x *bcd* embryos unless *fat2RNAi* knockdown is further introduced into this genetic background (Fig 5C-D; Fig S4D). A large percentage of these individuals showed abdominal patterning defects, with A4 being the most susceptible position (Fig 5C-D; Fig S4B). Interestingly, a similar distribution of defective abdominal segments is also seen in the small proportion of non-hatched *fat2RNAi* individuals (Fig 5C-D).

To understand if embryonic geometry predetermines the severity of phenotypic defects in individuals with *bcd* overexpression, we characterized the patterning outcomes using En expression in embryos with various *bcd* copy number and embryonic length. Individuals with 4x *bcd* (single insertion of tandem *bcd*), within the natural range of embryonic geometry, showed intact En expression. However, shorter embryos (<450 μm) frequently presented patterning defects, most commonly in the 6th En stripe (Fig 5E, top panel). Interestingly, this position corresponds to the A4 segment in the cuticle pattern. Comparatively, patterning defects become more pervasive in 6x *bcd* individuals when embryonic length reduced below 470 μm. The range of defective segments gradually expands from the 6th En stripe to both anterior and posterior regions with decreasing embryonic length (Fig 5E, middle panel and 5F). Further, increasing *bcd* dosage to 10x renders patterning processes exceedingly susceptible to reducing embryonic length. Defects are observed in comparatively shorter individuals within the natural range and recurringly the 6th En stripe is the most frequent breaking point in the patterning (Fig 5E, bottom panel).

The defective abdominal patterning that we observe here is an intuitive result, as both the posterization of gap gene boundaries due to increased *bcd* dosage and reduced embryonic length lead to reduced number of nuclei along the AP axis in the trunk region. When the number of nuclei falls short of the minimal requirement to fulfill all the different cell identities along the AP axis, certain cell fates become lost. It is surprising, however, that the position of lost cell fates is not stochastic, but originates at and expands from the A4 segment. This positional bias is also reflected in the segmentation gene pattern at the blastoderm stages. While the gap gene boundaries remain roughly at the same scaled positions across different geometry (Fig S4E), the absolute distance between neighboring gap gene expression peaks decreases in response to reduced embryonic length. This in turn changes the combinatorial inputs to activate downstream pair-rule genes, e.g. *even-skipped* (*eve*). Fig 5G-H illustrate that the expression peaks of Kni and Gt are brought into proximity with gradually reducing embryonic length. As Kni and Gt confine the boundaries of *eve* stripe 5 [34], the expression of this *eve* stripe is over-repressed (Fig 5H, asterisk). This results in the loss of correct cell fate at this position, corresponding to the future A4 segment. With further reduced embryonic length, a larger percentage of individuals fail to activate Kni and Gt in the trunk region (Fig 5G, green crosses; Fig 5H, arrowhead), leading to defects across a broader range.

## Discussion

Individuals of the same species often display a certain level of morphological and behavioral differences, such as in animal color patterns and human facial features [35,36]. This reflects inter-individual variation in genetic composition and life-history environmental exposure [37]. Such intraspecific individuality may have significant ecological and social impacts on the population [38]. Equally, these same genetic and environmental variations pose challenges to the fundamental developmental processes, as they are to generate invariant developmental outcomes. Multiple evidences suggest that organisms have evolved canalization mechanisms that render developmental processes insensitive to such sources of variation [2,39]. Early *Drosophila* embryonic patterning provides an excellent example of a canalized developmental process – the boundaries of segmentation gene expression remain highly reproducible amongst individuals in the face of heterozygous mutations [40,41], genetic variations [42] and temperature perturbations [28,43]. These studies suggest that mechanisms including epistasis, genotype-environment interactions and canalizing gene regulatory networks [44] work together to ensure precise patterning outputs.

In this study, we have identified embryonic geometry as an additional source of variation that patterning processes have evolved to buffer against. The geometry, or in other words, the aspect ratio of each ellipsoid-shaped embryo is determined during oogenesis, and this parameter varies by ±10% in the population of the wild-type strain OreR. The variable geometry in turn increases the variation in embryonic length given the natural range of embryonic size. Previous studies have shown that patterning outcomes are highly reproducible and remain scaled to embryonic length [42,45]. Correspondingly, we found that under decanalized conditions, either by depleting maternal *bcd* inputs or artificially increasing the *bcd* dosage, the patterning process loses its capacity to buffer embryonic length variations. Consequently, the length of an individual embryo predetermines its patterning outcomes. The predictive power becomes stronger when we artificially increase the variation of the embryonic geometry. The aspect ratio of the *fat2RNAi* embryos differs by ±30% while the average embryonic size is only slightly reduced by ~8 %. These results further support embryonic geometry as a major source of variation that accounts for interindividual phenotypic variation under decanalized conditions.

Both embryonic size and embryonic length are inheritable traits and therefore adaptive to artificial selection or environmental changes [42,46–48]. It will be interesting to understand if the aspect ratio of the embryo shape is also a genetically variable trait so that the population can be selected to produce progenies with a biased geometry. If this is the case, embryonic geometry may be involved in the complex interplay between environment, genetic components and developmental processes during the course of evolution. When a population confronts selection towards a new phenotypic optimum, for example, larger egg size due to decreasing temperature [47], such directional selection may result in decanalizing effects on the patterning processes [46,49]. Meanwhile, the naturally variable embryonic geometry – together with other sources of variation – generates a spectrum of patterning outcomes in different individuals. As a result, a different range of embryonic geometry will be favored and selected as they maintain the patterning outcomes of the parental lines. Conceivably, this may be one of the reasons why eggs of closely related Dipteran species differ not only in size [50,51] but also in geometry, and such geometrical differences can also be observed in different laboratory lines carrying different genetic background.

Our quantitative analysis of segmentation gene expression demonstrates how embryonic geometry affects individual patterning outcomes under two decanalizing conditions. In the case of the maternal *bcd* null mutant, we have shown that the signaling centers located at both poles of the embryo initiate the hierarchical gene expression along the AP axis in a non-scaled manner. This explains, mechanistically, how patterning processes incorporate information of the embryonic geometry to account for the final outputs. It remains unclear, however, in the case of increased *bcd* dosage, what determines the breaking point (the fourth abdominal segment) of the final pattern. One possibility is that the susceptibility of this position reflects the strength of the regulatory interactions between the segmentation genes [52]. Systematic comparisons among different *Drosophila* species have shown that the regulatory sequences of the segmentation genes are rapidly evolving and thus substantially diverged [53]. Interestingly, the spatio-temporal dynamics of the segmentation gene expression patterns are highly conserved between species, suggesting that the co-evolution of modular transcription binding sites compensate for each other to keep the patterning outcomes unchanged [53–55]. Such an inter-species canalization phenomenon is also observed among more distally related species within the sub-taxon Cyclorrhapha, which involved more dramatic rewiring of the regulatory network [56,57]. If the breaking point of patterning processes under decanalized conditions truly depends on the system parameters of the underlying network [58], we expect to see different susceptible points in different network structures. This can be tested by characterizing decanalizing phenotypes in related species.

In conclusion, embryonic geometry was identified as a source of variation in addition to environmental and genetic factors that predetermines phenotypic outcomes in mutant conditions. We think that embryonic or more generally, tissue geometry may play an important role in other decanalizing conditions by affecting patterning outputs, such as other segmentation gene mutants [14,15], or *in vitro* induction of patterning systems [59,60], both of which show significant inter-individual phenotypic variations. Our work highlights that care must be taken when taking a system out of its native environment – *e.g.* organoids – as the system boundaries affect the operation of signaling networks. Characterizing the influence of the geometrical parameters will help us to have a more complete understanding of decanalization, and in turn, canalizing phenomenon.

## Materials and methods

### Fly stocks and genetics

The bcd knockout allele (*bcd^KO^*) used in this study was generated by CRISPR-mediated insertion of a MiMIC cassette into the first intron of the *bcd* gene [19,20] The cuticle phenotype of *bcd^KO^* was compared to that of the classic *bcd^E1^* allele [61] To generate embryos with artificially reduced aspect ratio, we expressed RNA interference against the *fat2* gene using a maternal *traffic jam (tj)>Gal4* driver [62] Both *UAS>fat2RNAi* and *tj>Gal4* were either crossed to or recombined with the *bcd^KO^* allele, so that the females carrying all three alleles produce bcd null embryos with wide range of aspect ratio.

The *tandem bcd* construct was generated by replacing the eGFP sequence in the pCaSpeR4-*egfp-bcd* vector [26] by *bcd* protein coding sequence. First, the vector was digested with NheI and SphI to remove the eGFP. Next the *bcd* coding sequence was amplified by PCR from the vector using primer pairs 5’-cggagtgtttggggctagcaaagatggcgcaaccgccg-3’ and 5’-gttagtagctccgcttccattgaagcagtaggcaaactgcgagtt-3’.

Further the P2A self-cleaving peptide with the GSG linker was synthesized as oligo pairs 5’-tttgcctactgcttcaatggaagcggagctactaacttcagcctgctgaagcaggctggagacgtggaggagaaccctggacctgcatgcatggcgcaaccgc-3’ and 5’ggcggttgcgccagcatgcaggtccagggttctcctccacgtctccagcctgcttcagcaggctgaagttagtagctccgcttccattgaagcagtaggcaaa-3’.

These two fragments and the digested vector were then assembled using Gibson Assembly strategy (NEB). The final construct was injected (BestGene Inc.) and two insertions on 2nd (*tdBcd(II)*) and 3rd (*tdBcd(III)*) chromosome, respectively, were established and used for this study. Consequently, hetero- or homo-zygous *tdBcd(III)* females produce embryos with 4x and 6x of maternally loaded Bcd protein, respectively (compared to 2x*bcd* in the wild-type); and homozygous *tdBcd(II);tdBcd(III)* females generate 10x*bcd* embryos. Finally, females homozygous for *tdBcd(III)* which also carry *tj>Gal4* and *UAS>fat2RNAi* generate 6x*bcd* embryos with reduced aspect ratio.

Other fly lines used in this study include a laboratory OreR strain raised in 25°C (the wild-type control); *en>mCD8-GFP* (to visualize dynamic *en* expression pattern); *egfp-bcd* line (for quantification of Bcd gradient profile); *mat>eGFP; hb>LlamaTag* (gift from Hernan Garcia’s lab); *Df(3R)tll^g^* (BL#2599).

### Measurement of embryonic geometry

To compare the geometrical parameters between OreR, *bcd^KO^* and *fat2RNAi* populations, embryos were dechorionated and aligned laterally on an agar plate and imaged under a stereoscope (Nikon SMZ18). Images were then segmented to extract the embryo contour and fitted to elliptic shapes. The long and short axes of fitted ellipses were taken as the measurement of embryonic length and width, respectively. For each strain, more than 200 individuals were measured.

For confocal live imaging, embryonic length was measured as the longest distance between the anterior and posterior poles. For immunostained embryos, fixation results in an isotropic shrinkage of embryonic volume. To measure the geometrical parameters of the fixed embryos, we first carried out a linear fit between aspect ratio and embryonic length using stereoscope data. Further we measured the aspect ratio of each fixed embryo and estimate its embryonic length and width using the same linear fit equation.

### Immunostaining

Embryos at desired stages were dechorionated by household bleach and fixed in heptane saturated by 37% paraformaldehyde (PFA) for 1 hr. The vitelline membrane was subsequently manually removed. Prior to incubation with primary antibodies, embryos were blocked with 10% BSA in PBS. Antibodies used were guinea pig anti-Hb (1:2000), rabbit anti-Gt (1:800), guinea pig anti-Kr (1:800), guinea pig anti-Kni (1:800), guinea pig anti-Eve (1:800). Primary antibodies were detected with Alexa Fluor-labelled secondary antibodies (1:500; LifeTech). Embryos were co-stained with Phalloidin conjugated with Alexa Fluor for staging purpose or visualizing cephalic furrow position. Short incubation of Dapi dye was carried out during the last wash prior to mounting to visualize presyncytial nuclei. Embryos were mounted in AquaMount (PolySciences, Inc.) and imaged on a Zeiss LSM710 microscope with a C-Apochromat 40x/1.2 NA water-immersion objective. Hb, Gt, Kr, Kni and Eve antibodies were gifts from Johannes Jaeger.

### Cuticle preparation

Embryos of various genotypes were collected during the blastoderm stages and allowed to develop at 25°C until the end of embryogenesis. The embryos were then dechorionated, fixed, devitellinized and incubated into a mixture of Hoyer’s medium and Lactic acid in a 1:1 ratio at 65 °C between an imaging slide and a cover slip. For an exhaustive description of the method used see Alexandre (2008).

### Measurement of Bcd profile

For measurement of Bcd gradient profile, we followed the protocols detailed in Gregor *et al.* [4]. Embryos expressing eGFP-Bcd either with or without *fat2RNAi* were dechorionated and mounted laterally on a confocal microscope (Zeiss LSM710). The images were acquired at the midsagittal plane of embryos at early n.c. 14. Images acquired for different individuals were taken with identical microscope settings. For each image, nuclear centers along the dorsal edge of the embryo were manually selected and the corresponding circular area was used to compute the average fluorescent intensity. Nuclear intensity was then plotted against either absolute distance from the anterior or scaled AP position. To compare average profiles between control and *fat2RNAi* embryos, all nuclei from embryos either longer or shorter than 450 μm are binned in 50 bins along the scaled AP axis over which the mean and standard deviation were computed.

### Simulation of SDD model

We simulated using Matlab the SDD model in steady-state for morphogen concentration, *ρ*(*x,t*) at position *x* and time *t*:

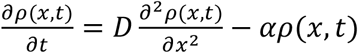

where *D* is the diffusion coefficient and α the degradation rate. We approximated the *Drosophila* embryo as a cylinder, with different lengths (500μm, 400μm) and radii (90μm, 130μm) for the wild-type and *fat2RNAi* embryos respectively. The effective radius in the simulation for *fat2RNAi* embryos is slightly larger than measured experimentally (~110μm), reflecting the rounder geometry in these embryos (hence the cylindrical approximation of embryo shape is less accurate). Using an elliptical geometry does not significantly alter these results (data not shown). We account for eGFP folding time (~50 minutes) [63], and the results plotted in Figure S2D are for the folded population of Bcd::eGFP.

### Gap gene boundary quantification

Confocal Z-stack images were Z-projected (maximum intensity) in Fiji (RRID:SCR_002285) for further analysis. The images of laterally oriented embryos were rotated so that the anterior is to the left and dorsal to the up. A line with the width of 100 pixels crossing the center of the embryo was drawn to abstract average intensity along the AP axis. The intensity profile was shown as a function of percent embryonic length (%EL). The boundary positions of gap gene expression domains were defined as the position (% EL) where the intensity corresponds to half maximum intensity value of the domain. To estimate the variability in the precision of boundary position, we performed bootstrapping using the Matlab function *bootstrp*. We performed 1000 simulation runs to infer the variability on the boundary precision. As the number of samples per boundary was small, we did not test for significant changes in the precision of boundary specification for a single boundary between wild-type and *bcd^KO^* embryos. However, pooling the data from the different boundaries, we observe that precision in all the measured boundaries decreases (*i.e.* the error increases) in *bcd^KO^* embryos. We calculated the p-value using the paired sample t-test across all boundaries.

### Quantification of maternal bcd transcripts

To compare the relative amount of maternally loaded *bcd* transcripts in different genotypes, we extracted total mRNA from presyncytial embryos (within 1h after egg deposition) generated by OreR, *fat2RNAi*, *tdBcd(III)* or *tdBcd(II);tdBcd(III)* females and reverse transcribed to cDNA. We performed qRT-PCR with *bcd-*specific primer pair using SYBR Green (Thermo Fisher) protocol and the housekeeping gene *rpl32* was used as internal reference. The relative *bcd* mRNA amount was normalized to that of OreR. Three independent measurements were carried out over which the mean and standard deviation was calculated.

## Acknowledgements

We thank Sally Horne-Badovinac and Hernan Garcia for fly lines. We thank Alexis Kerh and Jean-François Rupprecht for help with fly work and modeling of the Bcd gradient respectively. This work was supported by a National Research Foundation Singapore Fellowship awarded to T.E.S. (NRF2012NRF-NRFF001-094) and funding from the Mechanobiology Institute, National University of Singapore, Singapore.

## Author Contributions

AH and TES conceived and designed the project. AH performed all experiments. AH analyzed and quantified the data with assistance from TES. TES performed model fitting. Both authors wrote the manuscript.

## Supplementary Figures

**Fig S1.**
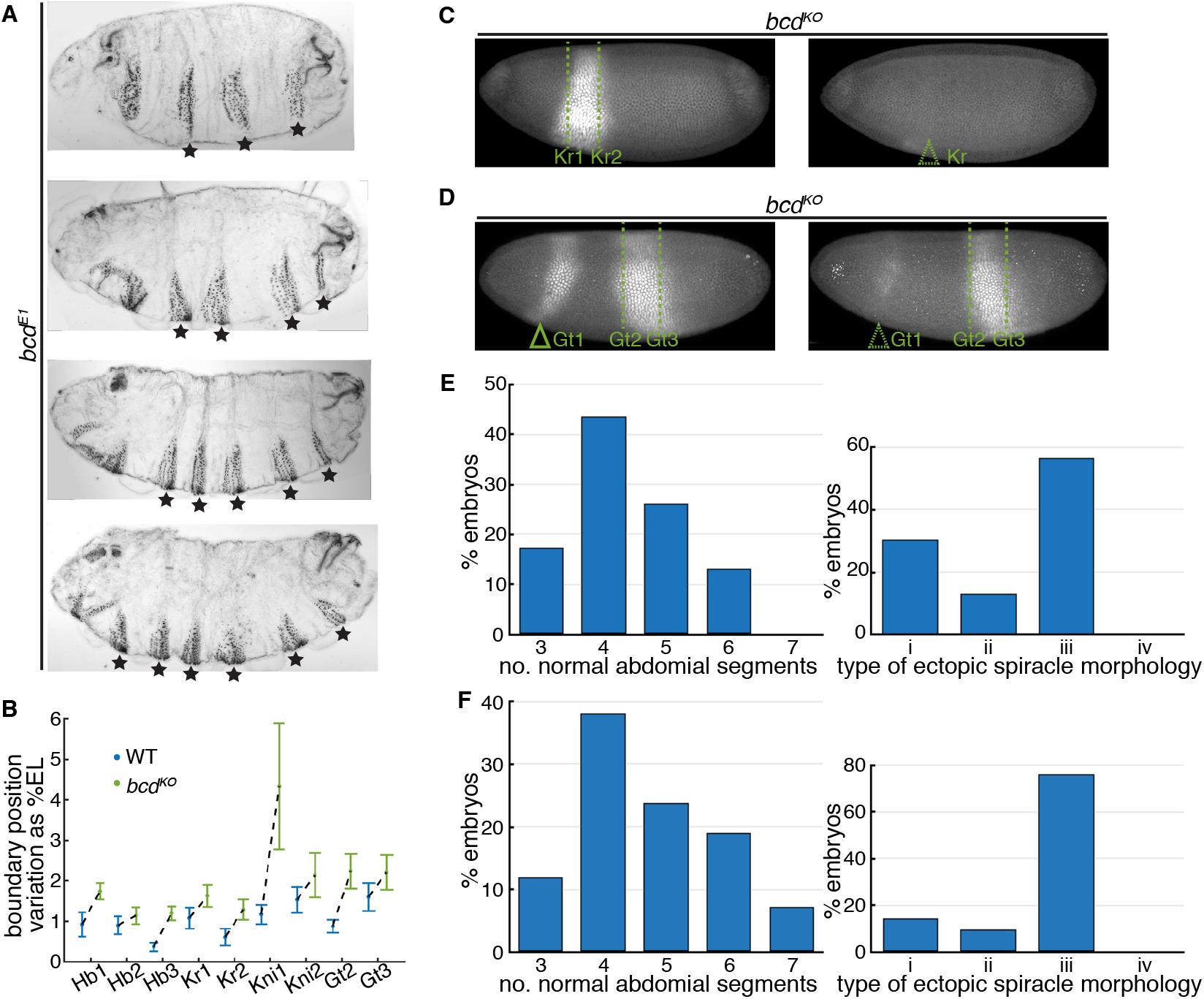
Phenotypic variation in *bed* null mutants. (A) Representative cuticle patterns of embryos with maternal *bcd^E1^* allele, showing different number of normal abdominal segments as indicated by stars. (B) Variation of gap gene boundary positions in wild type vs. *bcd^KO^* embryos. Error bars are computed by bootstrapping with data shown in Fig 1J-M. (C-D) Comparison of *bcd^KO^* individuals showing different expression patterns of Kr (C) and Gt (D). Solid triangle indicates anterior Gt domain and dashed triangles indicate failed activation of Kr or anterior Gt. (E-F) Phenotypic frequency of cuticle patterns and ectopic spiracles in individuals generated by a single *bcd^KO^* female crossed to a single male and raised in constant environments. E and F show two independent single-cross experiments. n = 23 (E) and n = 42 (F).

**Fig S2.**
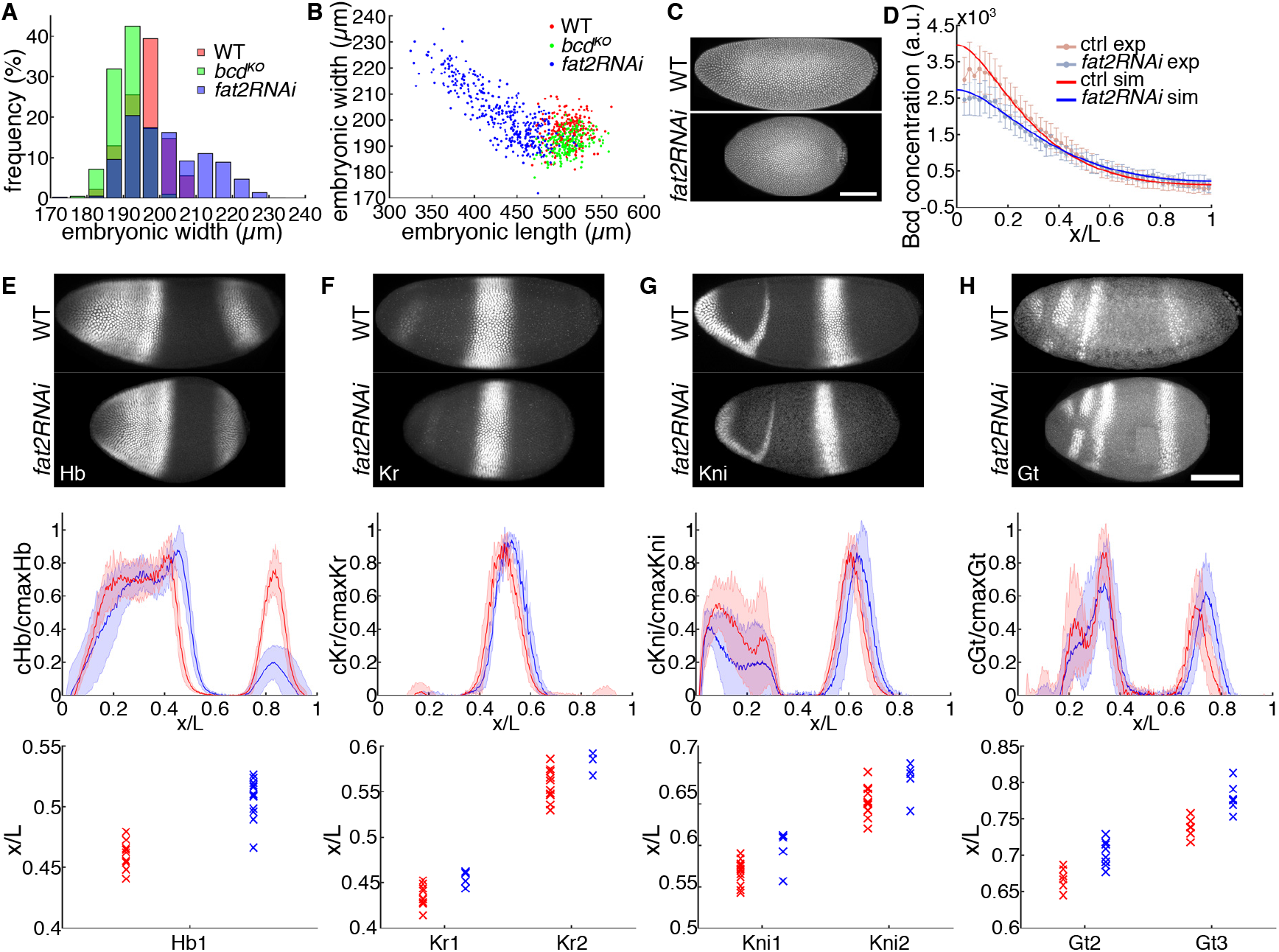
Effects of perturbed embryonic geometry on early patterning. (A) Distribution of embryonic width in wild type (n = 239), maternal *bcd^KO^* (n = 210) and *fat2RNAi* (n = 364) embryos. (B) Embryonic width plotted against embryonic length in three genotypes. (C) Max projection of Dapi staining in wild type and *fat2RNAi* embryos. (D) Bcd intensity profile along AP-axis for control (red points, n = 27) and *fat2 RNAi* embryos (blue points, n = 17), errorbars show s.d.. Equal uniform background subtracted from both profiles. Curves represent fitting to solution of steady-state SDD model along a cylinder, as described in methods. Identical diffusion and degradation parameters used for both fits. (E-H) Comparison of gap gene expression between wild type and *fat2RNAi* embryos. Profiles were normalized to the max intensity and the computed mean and s.d. were plotted against scaled AP length. Boundary positions of each individual is plotted to show the distribution in two genotypes (WT, red and *fat2RNAi*, blue). Scale bar, 100μm.

**Fig S3.**
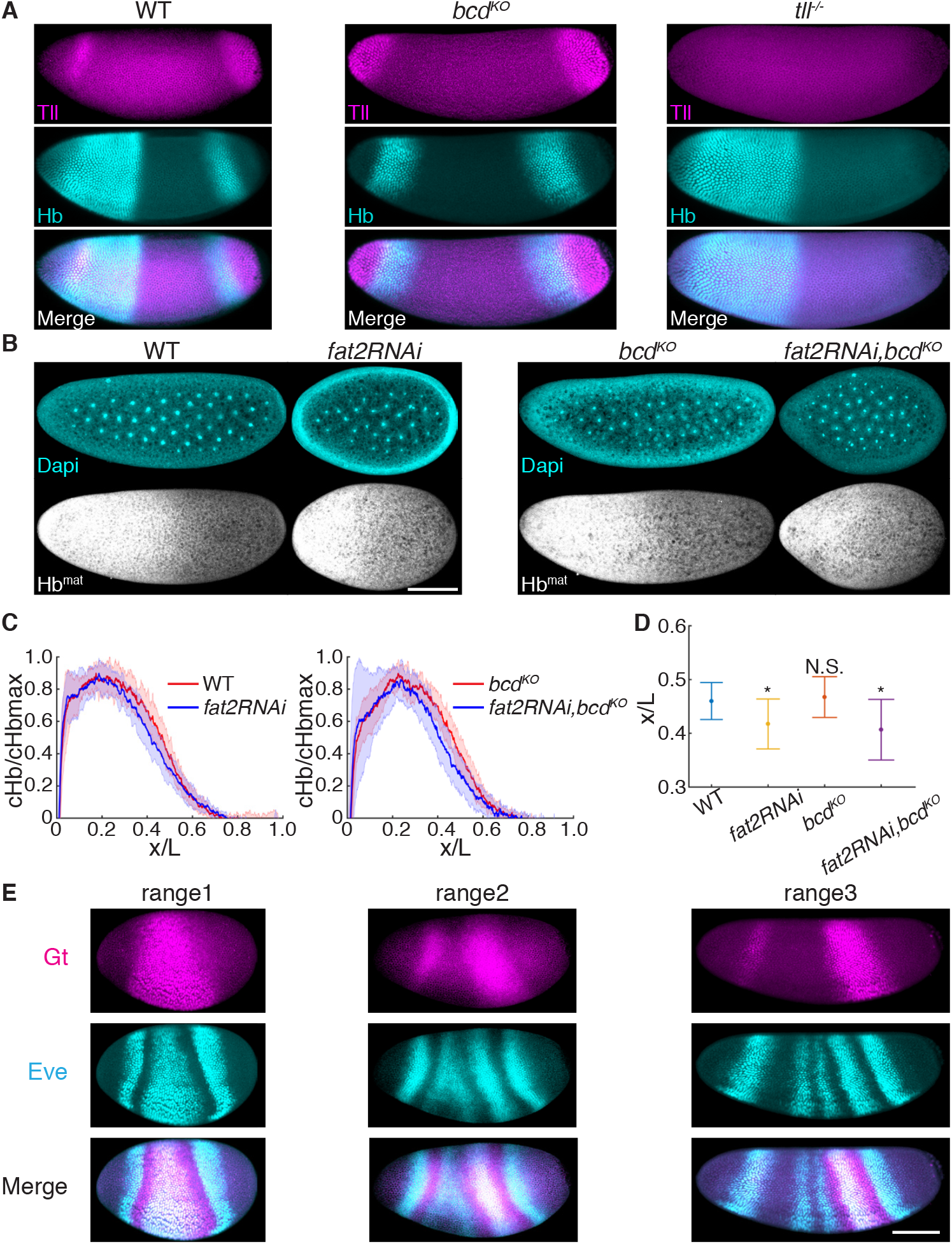
Positional information transfer in the absence of maternal *bed*. (A) Embryos of wild type (left), *bcd^KO^* (mid) and *tll* (right) fixed at the end of the blastoderm stage and stained for Tll (magenta) and Hb (cyan). The symmetric expression of Hb and Tll in *bcd^KO^* and the absence of posterior Hb stripes in *tll* embryos indicate that Tll is necessary for the activation of this Hb domain. (B) Embryos of wild type, *bcd^KO^*, *fat2RNAi*, and *fat2RNAi*, *bcd^KO^* fixed at presyncytial stages (around n.c. 8) and stained with Dapi and Hb antibody. The Hb intensity at these stages represent maternal Hb expression. (C) Mean and s.d. of normalized Hb profiles from four genotypes, respectively, were plotted vs. scaled AP position. (D) Mean and s.d. of maternal Hb boundary positions in four genotypes. *p<0.01. (E) Representative images of *bcd^KO^* embryos within different EL ranges (corresponding to Fig 4F) fixed at the end of blastoderm stage and stained for Gt (magenta) and Eve (cyan). Scale bar, 100μm.

**Fig S4.**
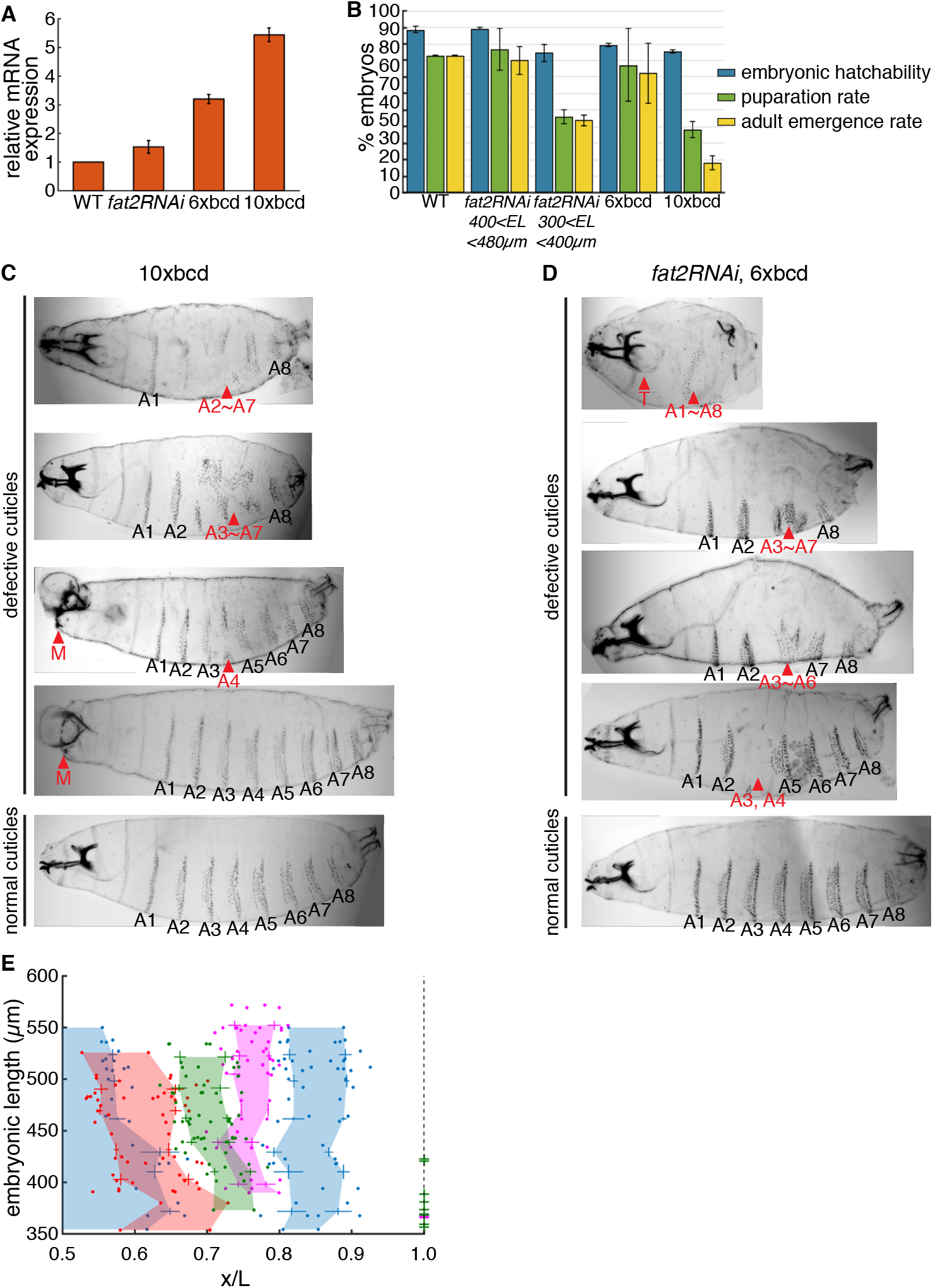
Embryonic patterning with maternal *bcd* overexpression. (A) Expression level of maternally loaded *bcd* mRNA in *fat2RNAi*, 6x*bcd* and 10x*bcd* relative to wild type. (B) Rate of embryonic hatching, puparation and adult emergence in wild type, *fat2RNAi* individuals longer or shorter than 400μm, 6x*bcd* and 10x*bcd* embryos. (C-D) Representative cuticle patterns of 10x*bcd* (C) and 6x*bcd* (D) embryos. Red arrowheads indicate defective regions. M, mouthparts and T, thoracic segments. (E) Embryonic length of 6x*bcd* individuals plotted against the scaled AP position of four gap gene boundaries (Hb, blue; Gt, magenta; Kr, red and Kni, green). Data from every 30μm EL interval were binned to compute mean and s.d. and the colored areas are generated by connecting mean values of different EL ranges. Dashed line indicates posterior boundary of individuals; green and magenta crosses overlapping the dashed line indicate individuals with corresponding EL not expressing Kni and Gt, respectively.

## Supplementary Videos

**Video 1. Engrailed expression in *bed^KO^* mutant individuals**

Live imaging of *bcd^KO^* embryogenesis from onset of gastrulation to the end of dorsal closure. Embryos express H2Av::mCh (red) and en>mCD8::GFP (green). Dots indicate En stripes and the numbers on top of the embryos indicate the AP length of each individual.

**Video 2. Embryogenesis of different geometry**

Wide field movies of wild type (top) and *fat2RNAi* (bottom) embryos from the onset of gastrulation until hatching.

**Video 3. Defective morphogenesis in late embryo development due to extreme embryonic geometry.**

Confocal movies of *fat2RNAi* embryos expressing *en*>mCD8::GFP from germband retraction to dorsal closure stages. Two embryos in the movie represent individuals with EL shorter (top) and longer (bottom) than 400μm. The dashed box indicates mismatch between opposing ectodermal tissue during dorsal closure. The numbers indicate the EL of each embryo.

